# Innovative beer-brewing of typical, old and healthy wheat varieties to boost their spreading

**DOI:** 10.1101/114157

**Authors:** Lorenzo Albanese, Rosaria Ciriminna, Francesco Meneguzzo, Mari Pagliaro

## Abstract

Certain typical old wheat varieties grown in the Mediterranean countries and elsewhere are renowned for their nutritional value due to higher concentration of polyphenols, especially flavonoids, as well as of important minerals, than modern varieties. However, the respective economical sustainability is constrained by reduced yield, increasing costs and stagnating or decreasing market prices. Moreover, under certain conditions, connected with the respective scale and local market volume, low-input, organically grown cereals can lead to lower environmental footprint. Brewing with a portion of such old grains can effectively boost their spreading, conditioned on its ability to supply a significant additional net income to farmers. Beer has become the worldwide most consumed alcoholic beverage and, although its market is dominated by few global players and standardized products, craft breweries have quickly spread out in most countries. Nevertheless, a scale issue has arisen about the economic sustainability of microbreweries, mainly due to high initial capital investment, energy costs, scale, and sometimes taxation. It is shown how breakthrough, cheaper, more efficient and profitable microbreweries can help solving the sustainability issues affecting old wheat varieties, thus boosting the respective spreading, lowering the environmental footprint of the cereal sector and contributing to improve the public health.

## 1. Introduction

Shifts in food production and consumption, involving cereals, have long been known to generate profound effects upon many categories of the environmental burden of the agricultural and food sector. A quite updated review of the present state of the art was offered by the special volume of the Journal of Cleaner Production entitled *“Towards eco-efficient agriculture and food systems”*, whose articles, along with many previous ones, were comprehensively reviewed by Sala and co-authors (Sala et al., 2017).

Italian authors investigated farming practices leading to minimization of the environmental burden from rainfed durum wheat cultivations under Mediterranean conditions (southern Italy), finding that a minimization of soil operations (no tillage), coupled with a suitable rotation with nitrogen-fixing legume crops, could increase the wheat yields and improve most categories of the environmental impact (Ali et al., 2015).

Lately, the same authors found that under optimized farming practices, including reduced nitrogen fertilizing and suitable rotation, the greenhouse gases (GHG) emissions could be lowered by over 70% both per unit area and per kilogram of produced wheat, with respect to conventional farming (Ali et al., 2017). Their results highlighted the wide room left for decisive improvements at the farming level and showed consistency with another study concerning a farmers’ cooperative in north-eastern Italy (Fantin et al., 2017).

In another paper of the above-mentioned special volume, Sonesson and co-authors introduced more advanced functional units than the simple mass-based one, *i.e.* “gram protein”, “gram digestible protein” and a more complex unit involving the digestible intake of the nine essential amino acids (Sonesson et al., 2016). Because of the new normalizations, the assessment of few key environmental impact categories for different food products, such as global warming potential, land use and freshwater ecotoxicity, clearly improved after accounting for the products’ nutritional value. However, neither the impact upon the public health, along with connected environmental and economic costs, nor additional food properties contributing to a healthy diet, such as antioxidant activity and substances, were considered.

In a further article of the same special volume, authored by Spanish scholars, the nutrition, dietary and public health issues were much more central to an original assessment of the impact of carbon-based food taxes on the mitigation of the carbon footprint by acting on the consumption side (García-Muros et al., 2017). Given the high food expenditure rate upon cereals (on average, 22.9%), along with the highest emissions attributable to meat and dairy products, at least in Spain the proposed tax leverage was assessed to trigger a shift of consumption towards cereals, vegetable and fruit and, it was argued, a healthier diet due to reduced intake of calories, proteins, and saturated fats, as well as increased intake of fibers.

However, admittedly the proposed taxation would be slightly regressive, although that flaw could be eased by adjusting the taxation itself, as well as quantitative economic health benefits associated with improving diets remained unaccounted. Moreover, cereals fell under a unique category without distinction, and further health benefits from possible increased uptake of specially healthy substances were not considered.

Some old wheat varieties are renowned for the respective nutraceutical properties, mostly due to greater concentration of powerful antioxidant compounds (AOC), mainly flavonoids, as well as of both soluble and insoluble fibers (Di Silvestro et al., 2016; Dinelli et al., 2011; Leoncini et al., 2012; Migliorini et al., 2016). Among those varieties, some of the most interesting are grown in Tuscany (Ghiselli et al., 2010).

Moreover, a substantial contribution to the nutritional quality of some old wheat varieties derives from the greater concentration of selected minerals fundamental of the human metabolism, among which zinc, which was found to be particularly abundant in few old landraces such as Verna (Migliorini et al., 2016). This mineral was recently indicated as a very important element reducing the risk of some cancer, cardiovascular and diabetic diseases, especially for populations with cereal-rich diets (Zyba et al., 2016).

Based on the widespread use of cereal products in the Mediterranean diet, the transition to healthier raw food ingredients could significantly improve the consumers’ health, reducing the associated social and sanitary costs, increasing labor productivity and possibly reducing the sanitary-related environmental burden. So far, to the best knowledge of the authors, possible benefits directly related to a shifting to healthier diets remained unaccounted in life cycle assessment (LCA) studies under both the economic and environmental sides.

About the environmental footprint of typical old wheat varieties, an LCA study of the bread supply chain in Europe revealed critical issues affecting low-input grain farming (Kulak et al., 2015). In particular, if low-input farming quite obviously lowers most parameters of the environmental footprint *per unit of cultivated area,* the same is far from obvious when coming to the impact *per unit quantity* of final product, largely due to the lower yields and consequent larger overall cultivated area, greater fuel and electricity consumption, larger infrastructure on the farming side. The scale issue adds to offset the benefits of low input of fertilizers and pesticides: the common practice of direct selling small volumes of products to end consumers, or to small shops, produces a far larger fuel consumption per unit product on the distribution and consumers’ side.

According to the same authors, while a relative spreading of typical old wheat varieties in Europe could be afforded due to plenty of supplies, with imports rising over domestic production, and declining agricultural land, the overcoming of the respective niche scale, leading to a relief of the transportation issue, would require a constraint on consumption of the overall products or their composition (Kulak et al., 2015).

Indeed, the economic sustainability of typical old wheat varieties has been limited by lower yields than modern ones, despite their gradual increase (Ghiselli et al., 2010). Moreover, in case of organic or low-input biodynamic crops, the yield gap further reduces, while few typical old wheat varieties grown under low-input farming practices, such as Verna, stand out for the respective nutraceutical properties (Dinelli et al., 2011).

Economic turmoil, increased price volatility, dependence of farming costs (fuel, nutrients and chemicals) upon the price of fossil fuels, constrain the space and increase the risk of investments in the overall agricultural and food sector, let alone the risk of shifting towards lower yield crops (Sommerville et al., 2014; Timmer, 2017). The wheat commodity price has indeed suffered a recent downfall, closely following the oil price and threatening farmers’ margins (IndexMundi, 2017).

In demographically depressed European countries, among which those bordering the Mediterranean, limited perspective exists for further expansion of the domestic cereals market, as well as the economic stagnation and the associated downward pressure on the demand side restrain from imposing any sort of food taxes with possible regressive consequences, such as the above-mentioned proposed ones (García-Muros et al., 2017).

The main purpose of this study is advancing a new purely market-based sustainability roadmap for the spreading of typical old wheat varieties, built upon cheaper, more efficient farm-level craft microbreweries using 100% or a lower fraction of such varieties of wheat in the respective beer recipes, able to contribute to the farmers’ net income. The technological foundations for such microbreweries were explained in a previous recent study by the authors (Albanese et al., 2017), to which the reader is referred for any detail. Anyway, few of its basic features will be recalled later on in Section 3.1.

The main underlying motivation for this study is the contribution to improve the public health in populations with cereals-rich diets. However, quantitatively accounting for the social, economic, and environmental benefits from a shift to healthier cereal-based diets remains outside the scope of this paper.

## 2. Beer, health and brewing old grains

### 2.1 Beer sector

With its very long history, beer-brewing practices have been long established (Pires and Brányik, 2015), its chemistry and biology are well known and documented (Bokulich and Bamforth, 2013), as well as comprehensive up-to-date publications covering the whole beer sector are increasingly available (Mosher and Trantham, 2017).

Nowadays, beer has become the worldwide most consumed alcoholic beverage, with about 200 billion liters per year, as well as with a strong identity mark (Ambrosi et al., 2014; Stack et al., 2016).

The beer sector has not been free to the great economic crisis erupted in 2008. Global annual beer consumption looks like to have peaked in 2013 with 197 billion liters, later falling to 193 billion liters in 2015 (Statista, 2015), while in the European Union (EU, 28 countries) the beer consumption is plateauing since 2010 just above 35 billion liters (The Brewers of Europe, 2016), down from almost 38 billion liters in 2007-2008 (The Brewers of Europe, 2010).

However, at least in Europe, the number of active breweries has almost doubled in only five years to about 7,500 in 2015, as well as exports have surged, leading to increased production despite the stagnating domestic consumption and signaling the strong vitality of the sector (The Brewers of Europe, 2016). Indeed, the trends to globalization and rising income experienced during few decades, and the consequent leveling of consumers’ choices, led to the strong rise of beer consumption in several emerging countries – Russia and China, the latter surpassing USA in 2003, are two striking examples – and a parallel slight decline in many traditional beer-consuming countries (Colen and Swinnen, 2016).

In Italy, a boom of craft breweries occurred in the last decade, so much that their number skyrocketed from about 100 in 2004 to around 750 in 2014, along with an unabated rising trend of consumption, with wine gradually replaced by beer as the preferred alcoholic beverage, as well as experimentation of new ingredients and local flavors (Fastigi et al., 2015). The crisis of the agricultural sector, with the percentage of agricultural area to total land area fallen from 70% in 1961 to 47% in 2011 (“FAO - Land Statistics,” 2017), the search for innovative jobs, the shift in consumers’ choices and the growing environmental and health awareness helped, In Italy, the emergence of the peculiar phenomenon of agricultural craft breweries, which reached the number of 72 in the year 2014 (Fastigi et al., 2016). The latter segment – agricultural craft breweries that, by law, must produce at least the majority of the cereal ingredients used in their beer production – is precisely the *vehicle* identified in this study to sustain the spread of typical old grains.

The economic sustainability of craft microbreweries has been in turn constrained by high capital costs (equipment, devices and facilities), as well as by operating costs such as deriving from taxation and energy consumption, the latter accounting for approximately 10% of the overall operative costs on the average of the beer-brewing sector and substantially higher for craft microbreweries (Kubule et al., 2016; Sturm et al., 2013). All the above, despite a growing share of consumers show their willingness to pay more for the best craft beers (Mascia et al., 2016).

As a further threat to sustainability of craft breweries, the above-mentioned economic stagnation prefigures a possible leveling of market volumes along with downward pressure on prices. Nevertheless, the niche market of microbreweries, as well as of brewpubs, although representing rather distinct business models (Moore et al., 2016), cannot be regarded as a mere ideological reaction to globalization in the name of “small is beautiful” or some sort of localism. Rather, the seed of the boom of craft brewing in the recent decades was implanted by the very same standardization of commercial beers, proceeded shoulder by shoulder with globalization and, more recently, with the aggressive merging and acquisition wave and the respective industry concentration (Madsen and Wu, 2016; Wells, 2016). So much that, as extensively reviewed by Madsen and Wu, inside the same beer style, most consumers cannot even be able to distinguish a brand from another, nor can they distinguish “premium brands” (e.g., premium lager beers) from their standard counterparts, the aggressive advertising campaigns being the ultimate and likely the only *raison d'*ê*tre* for such premium brands and the respective substantially higher prices (Madsen and Wu, 2016).

One of main reasons underlying the progressive merging and acquisition wave in the commercial beer sector, *i.e.* the high transportation costs of unit product, represents as well a strong point in favor of microbreweries and, even more, of brewpubs, conditioned upon the existence of a local market within reasonable distance, including sufficient total population with a high enough percentage of young males (Madsen and Wu, 2016; Moore et al., 2016). So much that, as pointed out by Wells, “high throughput breweries with process flow rather than batch production are more resource-efficient per unit of output than microbreweries, but some (and possibly all) of this is offset by greater environmental burdens in transport and storage” (Wells, 2016).

As analyzed by Madsen and Wu, and although limited to great brewery plants, while large economies of scale are still achieved in marketing and, to a lesser extent, in distribution, production costs show an apparent elasticity with revenues, or net sales. It means that economies of scale in production are viable only by means of multi-plant operations (Madsen and Wu, 2016). That’s interesting to the purpose of this study because, translating the result to microbreweries, i.e. to limited production volumes more suitable to the proposed brewing installation (Albanese et al., 2017), the envisaged savings in the production processes (energy, raw material, water) and possibly in plant’s capital cost could lead to a competitive advantage for small breweries. Furthermore, the same competitive advantage would allow microbreweries based upon typical old wheat varieties to spread and replicate while preserving small production volumes, therefore taking full advantage of the benefits from localization (Wells, 2016).

Under the guidance of the biggest brewery companies worldwide, showing large marketing cost shares, on average around 16%, moreover in turn leading to astonishing results in terms of net sales (Madsen and Wu, 2016), it is highly advisable that brewers of high specialty beers such as the ones based upon typical old wheats identify and exploit their own distinctive marketing message. Health properties of those specialty beers could be a viable one.

### 2.2 Beer, a healthy food?

Even brewed with standard ingredients – water, barley malt with the possible adjunct of modern cereal varieties, hops and yeasts – beer itself can be considered a healthy food, under certain assumptions.

Recently, a consensus study, led by an Italian team and carried out by means of an international expert panel, showed that a moderate beer consumption by healthy subjects can significantly lower the risk of developing cardiovascular and neurodegenerative diseases, mainly as a result of the assumption of specific polyphenols from malts, grains and hops (de Gaetano et al., 2016). Few years earlier, Piazzon and coauthors carried out an extensive study on commercial beers, finding out surprisingly high concentrations of valuable phenolic acids as well as antioxidant activity, especially in darker and stronger products, so much that they could conclude that “beer may represent an important part of the overall dietary intake of antioxidants, particularly in the form of phenolic acids” (Piazzon et al., 2010).

Specific and significant preventive action against Alzheimer disease from hops’ iso-α-acids, which are responsible for bitterness in beer, was recently argued along with its potentially deep social impact with rapidly ageing populations worldwide (Ano et al., 2017). Moreover, evidence has arisen for anti-carcinogenic action from hops’ prenil-flavonoids (Ferk et al., 2010; Karabin et al., 2015; Żołnierczyk et al., 2015), as well as from phenolic constituents extracted from both malt and hop (Ferk et al., 2010; Gerhäuser, 2005).

As well, beer was estimated as one of the richest sources of silicon in the diet, whereas that substance is known for its importance for the growth and development of bone and connective tissue (Casey and Bamforth, 2010).

The importance of the unveiled healthy effects of moderate beer consumption are of course connected with the extremely large consumption of that beverage, being an integral part of the lifestyles of whole populations worldwide. In this respect, any shift to other ingredients and adjuncts, in the event they can offer even superior nutritional properties, could lead to further widespread health benefits, let alone if such shift could boost the spreading of further healthier food products.

### 2.3 Brewing old grains

The use of cereals other than barley, including typical old wheats, or pseudo-cereals, although already common in traditional beers, has grown along with the spreading of craft breweries and the respective products, along with the rediscovery of traditional cereals.

Brewing a blend of malted barley (60%), malted commercial wheat (20%) and unmalted typical “Senatore Cappelli” wheat (20%), the latter grown in Sardinia region, Italy, resulted in a craft beer with higher polyphenol content and more balanced taste, in comparison with two other industrial beers made up of 50% malted barley and 50% malted commercial wheat (Mascia et al., 2014).

Brewing with up to 40% unmalted oat or sorghum was shown not only to be technically feasible, but also leading sometimes to better flavor and aroma in comparison with all-barley malt beers, despite the use of exogenous enzymes could be necessary especially in conjunction with oat (Schnitzenbaumer and Arendt, 2014).

Brewing with 100% unmalted grains was recently proven to be technically feasible by means of a commercial enzymes blend, finding that the resulting beers lacked in free amino-nitrogen (FAN), alcohol and color in comparison to all-malt beers, with unmalted wheat outperforming unmalted barley, oat and rye (Zhuang et al., 2016). Moreover, ample room is left for brewing optimization of both cereals such as wheat, and gluten-free pseudo-cereals such as buckwheat, quinoa and amaranth, for example by means of alkaline steeping to improve the respective malting processes (de Meo et al., 2011).

The growing market of gluten-free food represented a strong motivation to the brewing of pseudo-cereals and alternative gluten-free cereals such as rice, sorghum and maize, although with uncertain results on the side of sensorial features (Hager et al., 2014; Mayer et al., 2016). Ancient einkorn wheat *(Triticum monococcum),* which is also suitable for organic growing, has been investigated by Hungarian authors as a source of healthy AOC in beer brewing (Fogarasi et al., 2015). They found that both unmalted and malted einkorn enjoyed higher antioxidant activity and total polyphenol content than modern wheat varieties, comparable to unmalted barley and slightly lower than barley malt, allowing their conclusion that einkorn brewing could add beers some healthcare potential. Their results were confirmed by German authors who, investigating lipophilic antioxidant contents of wheat species, found that all the considered 15 einkorn varieties were ranked among the top 20 high-antioxidant varieties of all species (Ziegler et al., 2016).

Einkorn beers, along with emmer (*Triticum dicoccum*) and spelt (*Triticum spelta*) beers, are indeed produced and available on the market, especially in Italy and Germany, along with a plenty of other related products, from flour to biscuits and bread (Cooper, 2015). According to the latter author, emmer grain is particularly rich in fibers, while einkorn wheat – also especially suitable for growth over poor soils where modern wheat varieties fail – does contain gluten but its gliadin protein may be less toxic to sufferers of celiac disease in comparison to modern wheat. In the same study, breads from the ancient Kamut^®^ khorasan wheat – another emerging ingredient of beers – were mentioned for their higher antioxidant activities than whole grain modern durum wheat breads, especially after sourdough fermentation.

In a very recent study, goji berries were added to standard ale type beers at different times during the brewing process, finding that their addition just after lautering yielded the best results in terms of both total polyphenolics content, antioxidant activity, taste and appearance (Ducruet et al., 2017). The same authors pointed out the beneficial effect of a moderate beer consumption on the human immune system, strongly correlated with the content of the most valuable AOC. In summary, the conclusion can be made that certain typical old wheat varieties are especially rich in flavonoids and other phenolic compounds endowed with higher antioxidant activity than modern varieties. As well, beer itself produces significant, although variable, antioxidant activity due to extracts from malts, grains and hops, with recognized healthy effects under moderate consumption habits.

### 2.4 Healthier diets matter

The far-reaching relevance of the above stems from the fact that strengthening the human immune system by means of increasing the regular dietary intake of valuable AOC appears as a viable and effective way to improving public health along with easing sanitary-related costs and environmental burdens, especially in ageing populations typical of developed countries. Shifting diets towards higher AOC content by means of the spreading and consumption of typical old wheat varieties would ease as well the environmental burden of the agricultural sector. Among other emerging health threats on the global scale, spreading of antibiotic-resistant genes (ARG), often hosted in common pathogen microorganisms such as *Escherichia Coli,* has become an urgent health issue as well as a threat to economic stability (Ferro et al., 2016). Tackling such issue is challenging and multifaceted. Downstream, the main carrier of ARGs, *i.e.* wastewater, has to be effectively disinfected (Ciriminna et al., 2016; Zhang et al., 2016), as well as effective clinical countermeasures have to be undertaken to treat patients. Upstream, medical prescription and consequent release of pharmaceutical antibiotics should be kept as low as possible.

At least the latter two measures can significantly benefit from a widespread strengthening of the human immune system.

Effectivity of dietary intake of antioxidants to counter oxidative stress to the immune system and the related depression, normally increasing with age, has been known since many decades, as well as proven both *in vitro* and *in vivo* (De la Fuente, 2002). According to the same author, “antioxidants, … do not exert an indiscriminate stimulating effect on the immune cell function, but instead they are homeostatic factors. Thus, since the immune system is a health indicator and longevity predictor, the protection of this system by antioxidant diet supplementation may be useful for health preservation in the aged”.

Free from harmful side effects, adequate intake of natural AOC from foodstuff – mainly flavonoids and other phenolic compounds – was identified as a cheap and suitable means to counter both acute and chronic inflammations and prevent – and sometimes help treating – consequent acute and chronic illnesses such as neurodegenerative diseases, cancer, diabetes, rheumatoid arthritis, inflammatory bowel disease and cardiovascular diseases (Arulselvan et al., 2016; Pérez-Cano and Castell, 2016; Vezza et al., 2016).

In a recent comprehensive review, including *in vitro* and cellular investigations, Del Cornò and co-authors found that, because of their strong antioxidant power, the potential biological activity of many dietary polyphenols on cells of critical importance to the immune response, such as dendritic cells, can prevent and attenuate inflammation, as well as promote immunoactivation with consequent health benefits. So much that, in their own words, “the use of bioactive food compounds at pharmacologic doses is emerging as a preventive and/or therapeutic approach to target metabolic dysregulations occurring in aging, obesity-related chronic diseases, and cancer” (del Cornò et al., 2016). Nevertheless, quantitative information on doses for possible functional diets, exact regulatory mechanisms of the immune system including, as well as information on the qualitative/quantitative composition of food items are still scant and deserve further research.

## 3 Materials and methods

### 3.1 Brewing units

The same beer-brewing unit powered by controlled hydrodynamic cavitation (HC), with total volume capacity around 230 L, performing the mashing and hopping stages and comprehensively described in a previous study by the authors (Albanese et al., 2017), was applied to investigate the technical feasibility and the results of brewing with different fractions of typical old wheats. Physics and chemistry of controlled HC in liquid media were introduced in another recent study by the authors (Ciriminna et al., 2017).

HC-assisted beer-brewing at the real-scale experiments allowed finding a potential for successful industrial developments, along with significant advantages and no apparent drawbacks (Albanese et al., 2017). Dramatic reduction of saccharification temperature, increased and accelerated peak starch extraction, significant reduction of operational time after traditional stages such as dry milling and boiling are made unessential, represent the most important benefits, along with relevant energy saving, shorter cleaning time, volumetric heating which prevents caramelization and overall simplification of both structural setup and operational management of brewing processes. No cavitational damage to the equipment arose, as well as no wort and beer oxidation.

Figure 1 shows the experimental installation, described in detail in section 2.1 of the previous study by the authors (Albanese et al., 2017), along with the calculation, meaning and limitations of the basic HC metric, *i.e.* the Cavitation Number (CN), only sufficing here to recall the reference to the description of the cavitation reactor in the form of a Venturi tube (Albanese et al., 2015). All the tests ran in brew-in-the-bag (BIAB) mode, using the malts caging vessel shown in Figure 1. The same technological unit proved capable of reducing the gluten content in 100% barley malt wort and beers (Albanese et al., 2016).

**Figure 1.**
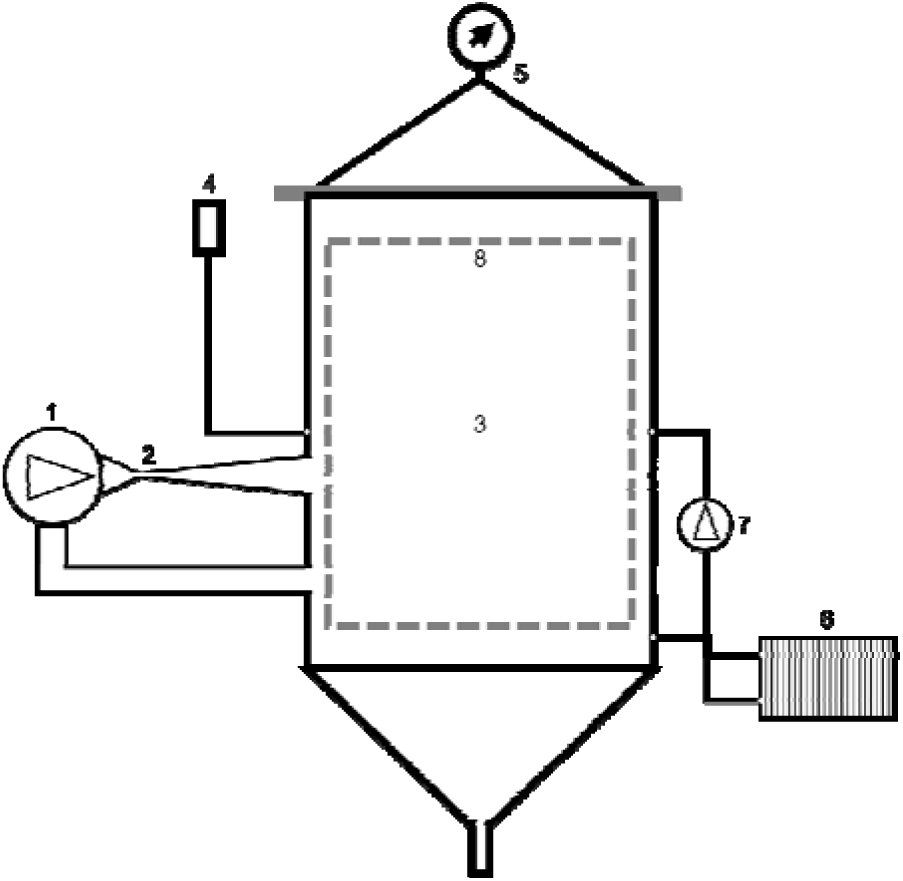
Simplified scheme of the experimental HC-based installation. 1 – centrifugal pump, 2 – HC reactor, 3 – main vessel, 4 – pressure release valve, 5 – cover and pressure gauge, 6 – heat exchanger, 7 – circulation pump, 8 – malts caging vessel. Other components are commonly used in state-of-the-art hydraulic constructions.

For comparison purposes, parallel brewing tests were performed by means of a conventional Braumeister (Ofterdingen, Germany) model 50 L brewer, equipped with a cooling serpentine (model Speidel stainless steel wort chiller for 50-liter Braumeister) and fully automatic brewing control (temperature, time and recirculation pumps).

### 3.2 Measurement instruments and methods

Along with thermometer and manometer sensors onboard the main production unit, few specialized off-line instruments were used to measure the chemical and physiological properties of wort and beer, described in full detail in a previous study by the authors (Albanese et al., 2017). Few of those instruments are described herein below.

Physico-chemical and physiological parameters of finished beers were measured by means of a 6-channel photometric device (CDR, Firenze, Italy, model BeerLab Touch). In particular, alcohol content (0-10% in volume, resolution 0.1%), color on the European Brewery Convention (EBC) scale (1 to 100, resolution 1) and on the Standard Reference Method (SRM) scale (0.5 to 50, resolution 0.1). All reagents were of analytical grade.

The power and electricity consumed by the three-phase centrifugal pump used in the HC device shown in Figure 1 were measured by means of a three-phase digital power meter (IME, Milan, Italy, model D4-Pd, power resolution 1 W, energy resolution 10 Wh, accuracy according to the norm EN/IEC 62053-21, class 1). The electricity consumed by the Braumeister model 50-liters conventional brewing device was measured by means of a mono-phase digital energy meter (PeakTech, Ahrensburg, Germany, model 9035, power resolution 1 W, energy resolution 100 Wh, accuracy 0.5%).

### 3.3 Production tests and brewing ingredients

Table 1 summarizes few basic features and results of the performed brewing tests. One set of tests (HC40 and B40) involved the use of Pale malted barley and a mix of raw unmalted grains from typical old wheat varieties, in the proportion 59% to 41%, respectively. The overall quantity of grains used in test HC40 was about 9% less than in test B40, in terms of proportion to the respective wort volumes.

**Table 1.** Beer production tests, ingredients and conditions.

| Test ID | Production unit<sup>a)</sup> | Net volume (L) | Grains<sup>b)</sup> | Exogenous enzymes<sup>c)</sup> | Hops<sup>d)</sup> | Beer color<sup>e)</sup><br>(EBC / SRM) | Wort SG<sup>f)</sup><br>/<br>Alcohol<sup>g)</sup> |
| --- | --- | --- | --- | --- | --- | --- | --- |
| HC40 | HC | 170 | Pale barley malt 21 kg<br>Old wheats mix 15 kg | Absent | Cascade 80 g | 16 / 7.1 | 1.039<br>/<br>5.3%vol |
| B40 | B | 50 | Pale barley malt 6.8 kg<br>Old wheats mix 4.8 kg | Absent | Cascade 26 g | 16 / 7.3 | 1.048<br>/<br>5.8%vol |
| HC100 | HC | 170 | Andriolo wheat 34 kg | CHILL 12 ml<br>PENTA 8 ml<br>BG SUPER 31 ml<br>AMYL 46 ml | Hersbrucker 0.3 kg | 4 / 2.1 | 1.017<br>/<br>2.6%vol |
| B100-1 | B | 50 | Andriolo wheat 11 kg | CHILL 4 ml<br>PENTA 2.5 ml<br>BG SUPER 10 ml<br>AMYL 15 ml | Hersbrucker 0.1 kg | 4 / 2.2 | 1.014<br>/<br>2.1%vol |
| B100-2 | B | 50 | Andriolo wheat 11 kg | CHILL 8 ml<br>PENTA 5 ml<br>BG SUPER 20 ml<br>AMYL 30 ml | Hersbrucker 0.1 kg | 4 / 2.0 | 1.012<br>/<br>1.8%vol |

a) HC = installation shown in Figure 1. B = Braumeister model 50-liters.
b) Grains were added from scratch in all tests. Wheat grains were raw and unmalted.
c) Exogenous enzymes were added from scratch in all tests.
d) Hops cavitating in any test.
e) Beer color was measured 15±2 days after bottling.
f) Wort Specific Gravity (SG) was measured in the wort before fermentation.
g) Alcohol content was measured 15±2 days after bottling.

The old wheats mix, supplied by Floriddia organic farm (Tuscany, Italy), included the varieties Andriolo, Frassineto, Gentil Rosso, Iervicella, Inallettabile, Maiorca, Sieve, Solina and Verna, approximately in the same proportion. The red varieties Andriolo, Gentil Rosso and Verna were attributed the highest antioxidant properties due to the respective free soluble and bound polyphenol and flavonoid contents (Di Silvestro et al., 2016; Ghiselli et al., 2016; Migliorini et al., 2016). Pelletized Cascade hops were used in those tests.

The other set of tests (HC100, B100-1 and B100-2) involved the use of 100% raw unmalted grains from the Andriolo old wheat variety, supplied by La Capannaccia farm (Tuscany, Italy). Again, the overall quantity of grains used in test HC100 was about 9% less than in tests B100-1 and B100-2, in terms of proportion to the respective wort volumes.

A blend of exogenous liquid enzymes, produced by Erbslöh Geisenheim AG, Germany, was used to perform saccharification. Namely, Beerzym CHILL (from the latex of papaya melons, increasing the degree of protein modification in brewing mashes), Beerzym PENTA (from a from a specially selected strain of *Trichoderma spec.,* degrading pentosan and glucan in original wort), Beerzym BG SUPER (from a specially selected strain of *Penicillium funiculosum,* degrading glucan in brewing mashes), and Beerzym AMYL (from a specially selected strain of *Bacillus subtilis,* liquefying starch in brewing with malt and raw grain portions). The above-mentioned exogenous enzymes were pitched in the same proportion to the respective wort volumes in tests HC100 and B100-1, while the respective quantities were doubled in test B100-2 in order to assess the sensitivity of results to the enzymes’ dosage.

Pelletized Hersbrucker hops were used in tests HC100, B100-1 and B100-2. Fermentation was activated in all tests by means of the dry yeast strain Safbrew WB-06, working at temperatures between 18°C and 24°C and tolerating a maximum alcohol content of 10.4%vol, always in the proportion of 50 g per 100 L of fermenting wort.

No simple sugar was added during the mashing stage in any test, while white sugar was added to the fermented wort before bottling and maturation, in the proportion of 0.85 kg per 100 L of fermenting wort in tests HC40 and B40, and in the proportion of 1.1 kg per 100 L of fermenting wort in tests HC100, B100-1 and B100-2.

## 4 Results

### 4.1 Brewing with 40% typical old wheat varieties

Figure 2 shows the basic features of the HC40 (a) and B40 (b) brewing processes. Mashing-out (extraction of malt and unmalted grains) was performed after saccharification, when the specific gravity (SG) values started plateauing. Despite a little more specific energy, in unit kWh/hL, was consumed in the HC40 test, the respective processes were fairly similar up to the boiling temperature. Subsequently, the long-lasting (80 min) boiling stage performed in the B40 test – which was unnecessary with the HC-assisted brewing device – caused a lower overall specific energy consumption in the HC40 test (23.4 kWh/hL) than in the B40 test (26.8 kWh/hL), *i.e.* an energy saving around 13%.

**Figure 2.**
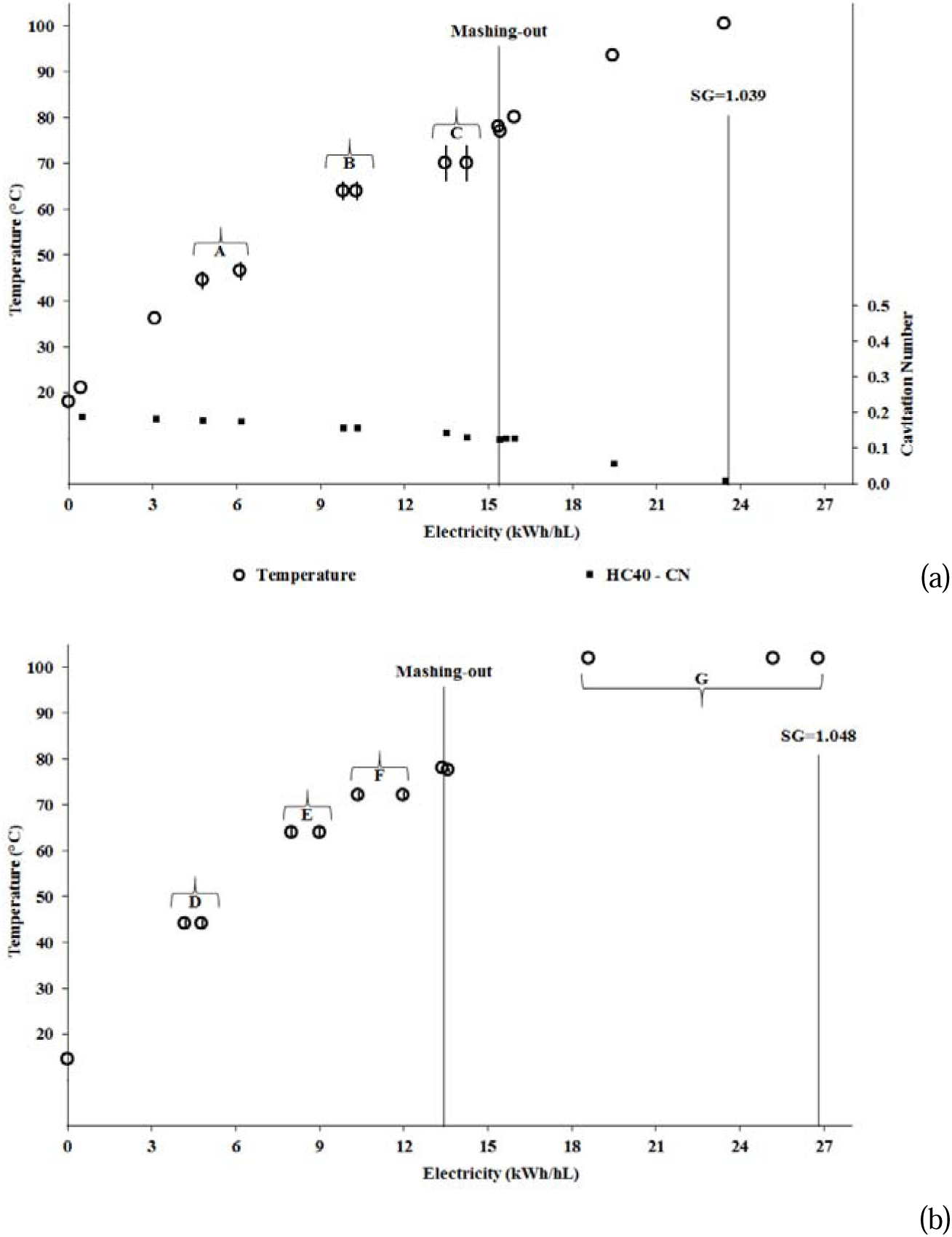
Temperature with respective uncertainties and cavitation number for test HC100 (a), and temperature with respective uncertainties for test B40 (b), both reproduced as a function of specific consumed electricity. Time of mashing-out, as well as value of specific gravity (SG) and its measurement time are indicated. Letters in the charts highlight the nearly isothermal steps: for test HC100, A (45.5±3°C, 53 min), B (64±2°C, 31 min), C (70±4°C, 46 min), for test B40, D (44±1°C, 45 min), E (64±1°C, 30 min), F (72±1°C, 45 min), G (boiling, 102°C, 80 min).

It should be noted that the BIAB set-up for the test HC40 was far from optimal, explaining why the energy saving was substantially lower than the figure – over 30% – found in the previous experiments with fully cavitating malts (Albanese et al., 2017).

Both the SG and the resulting alcohol content (Table 1) were a little lower in the test HC40, because of both the marginally lower proportion of grains (about 9% less than in test B40) and the suboptimal extraction of starch and enzymes with the BIAB set-up for the HC40 test.

Color data in Table 1 suggest similar Weissbier-style beers were produced (Priest and Stewart, 2006). Foamability for both beers were quite similar, while the subjectively assessed sensorial features suggested a remarkable superiority for the beer resulting from test HC40, not only in comparison with the beer resulting from test B40, but also across all the beers produced by means of the same HC-assisted brewing device, including those considered in the authors’ previous study (Albanese et al., 2017).

### 4.2 Brewing with 100% typical old wheat varieties

Figure 3 shows the basic features of the HC100 (a) and B100-1 (b) brewing processes. Mashing-out (extraction of unmalted grains) was performed after saccharification, when the SG values started plateauing. Starch conversion to fermentable sugars, which was carried out by means of the blend of exogenous liquid enzymes listed in Table 1, was surprisingly significantly higher in test HC100 than in test B100-1, especially with regards to energy consumption up to mashing-out (11 KWh/hL vs >18 kWh/hL). Moreover, also the SG observed in test HC100 was higher than in test B100-1 (1.017 vs 1.014), as well as the resulting alcohol content (2.6%vol vs 2.1%vol), despite the marginally lower proportion of grains (about 9% less in test HC100 than in test B100-1). Color data in Table 1 suggest Berlin Weisse-style beers were produced (Priest and Stewart, 2006).

As a consequence of the above, as well as of the absence of the boiling stage in test HC100, the overall specific energy consumption for the latter test (16.4 kWh/hL) was far lower than for test B100-1 (30.9 kWh/hL), translating into an energy saving around 47%.

A likely explanation of the latter result can be found in the increase of the mass transfer between exogenous enzymes and starch, as well as in the increase of starch gelatinization produced by HC processes (Albanese et al., 2017). The same results agrees with past experiences, such as dealing with HC-assisted ethanol production from dry mill corn using other blends of exogenous liquid enzymes (Ramirez-Cadavid et al., 2015). In the latter study, it was argued that HC-assisted processes could result in a diminished dosage of exogenous enzymes with equal results (*i.e.,* yield of fermentable sugars and ethanol itself). Similarly, from the above-described results one can argue that the dosage of exogenous enzymes, needed to brew 100% raw unmalted old wheat varieties in a HC-assisted device, can be substantially reduced in comparison with conventional brewing devices.

**Figure 3.**
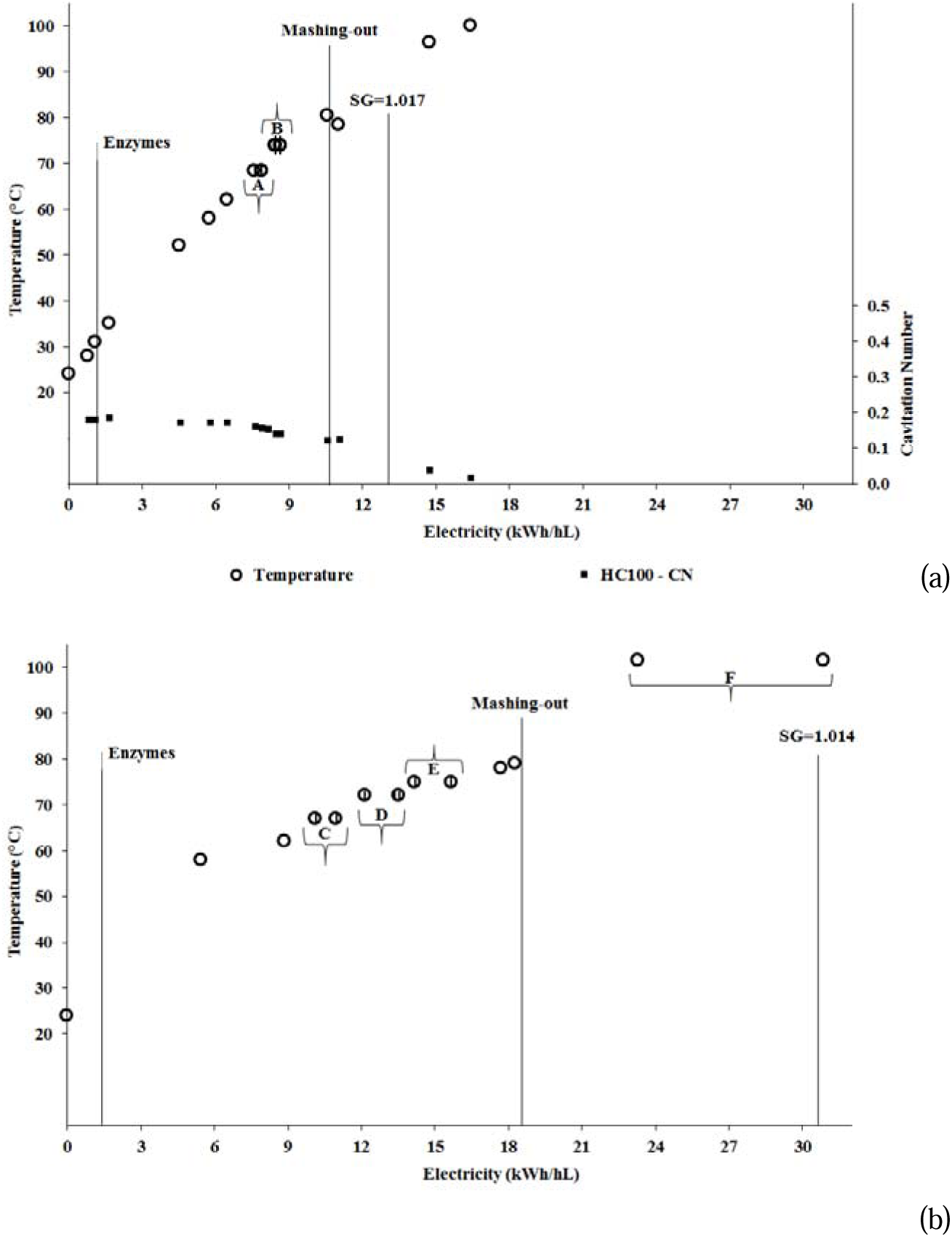
Same as Figure 2, for tests HC100 (a) and B100-1 (b). Letters in the charts highlight the nearly isothermal steps: for test HC100, A (68.5±1.5°C, 34 min), B (74±2°C, 76 min), for test B100-1, C (67±1°C, 21 min), D (72±1°C, 35 min), E (75±1°C, 37 min), F (boiling, 101.5°C, 71 min).

As shown in Table 1, the use of double dosage of exogenous enzymes in the conventional test B100-2 failed to provide better results in terms of SG and resulting alcohol content, further suggesting that the enzymes’ dosage could even be reduced with equal results, especially in the case of HC-assisted brewing, which is recommended as the subject of further research.

Subjective assessment of sensory features suggested that the products of tests HC100 and B100- 1 tasted as real beers, with the former one slightly preferred over the other one.

Overall, despite flaws such as too light color, too low alcohol content, and reduced free amino-nitrogen, as recently pointed out in recent studies (Pires and Brányik, 2015; Zhuang et al., 2016), as well as legal limits to the proportion of unmalted cereals in brewing recipes (Pires and Bränyik, 2015), brewing with 100% raw unmalted old wheat varieties is not only technically feasible but also potentially attractive.

Figure 4 shows a visual comparison between the resulting beers from tests HC40 and HC100, highlighting the apparent color difference (7.1 and 2.1 on the SRM scale, respectively), as well as the comparable and substantial foamability. Moreover, the pictures were taken about 3 months (HC40) and 4.5 months (HC100) after the respective brewing processes, suggesting as well an excellent foam stability, in agreement with previous results achieved with raw unmalted barley (Steiner et al., 2012).

**Figure 4.**
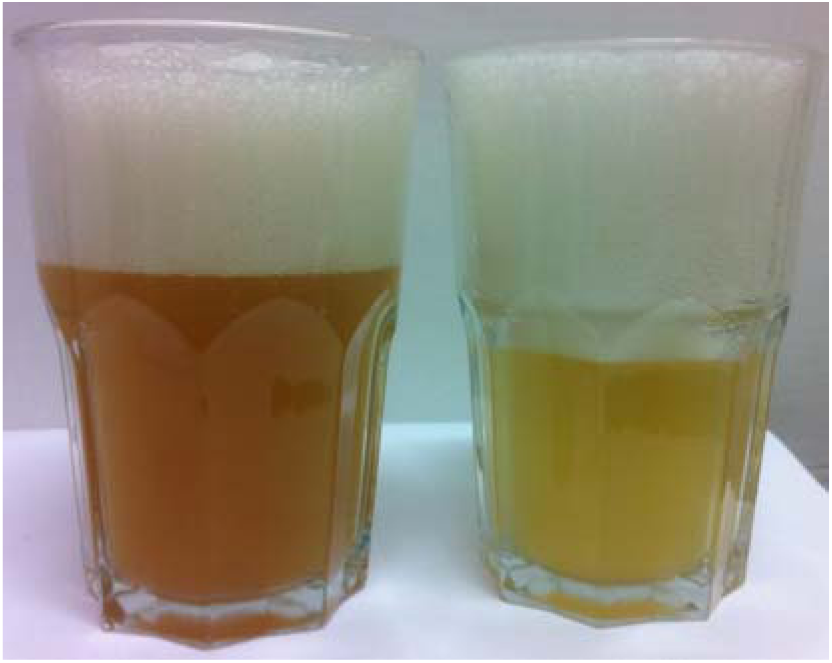
Beer samples produced from test HC40 *(left)* and test HC100 *(right).*

## 5 Discussion

Results shown in Section 4 confirm the technical feasibility of brewing beer with a proportion (40%) of raw unmalted old wheat grains, where saccharification enzymes are supplied by barley malt, as well as the same feasibility with 100% old wheat grains, using a blend of commercial exogenous liquid enzymes. Given the paucity of studies dealing with brewing 100% raw unmalted wheat (Zhuang et al., 2016), the latter appears as a fairly original result of this study. Moreover, such technical feasibility is shown by the novel HC-assisted brewing device and method too (Albanese et al., 2017), with distinct advantages in terms of energy efficiency, operation time, concentration of exogenous enzymes and even quality of end products.

In particular, despite the suboptimal BIAB set-up used for the HC-assisted brewing process, the overall specific energy consumption for test HC100 was 47% lower than in test B100-1, the latter performed with a conventional device, likely most attributable to the cavitational-induced increased mass transfer between exogenous enzymes and starch, and representing a quite original finding of this study in the field of beer-brewing.

HC-assisted brewing of 100% raw unmalted old wheat grains shows the greatest benefits over conventional techniques within the BIAB set-up, even if much work remains to be done in order to optimize the processes and fix the flaws of the resulting beers. Breweries, especially agricultural craft ones, deciding to use 100% raw unmalted grains, would gain further competitive advantage not having to resort to malt import from few large suppliers (Fastigi et al., 2016), or build their own micro malt houses with the consequent capital costs and environmental impacts (Steiner et al., 2012). In the latter study, advantages and disadvantages of brewing raw unmalted barley are listed and commented, and some of the disadvantages could be fixed with HC-assisted brewing in full-cavitation mode, which is recommended as a subject of further research.

On the other hand, it can be envisaged – but will have to be proven experimentally in future research – that in full-cavitating mode *(i.e.,* cavitating grains), the already excellent beer produced with 40% raw unmalted old grains will fully enjoy the efficiency gains achieved with brewing 100% barley malt (Albanese et al., 2017), thus offering breweries a wide flexibility in deciding their brewing recipes.

Figure 5 shows a simplified scheme of the vision developed in this study, along with the main logical connections, converging towards the widespread health and environmental benefits brought by the spreading of healthier typical old wheat varieties.

**Figure 5.**
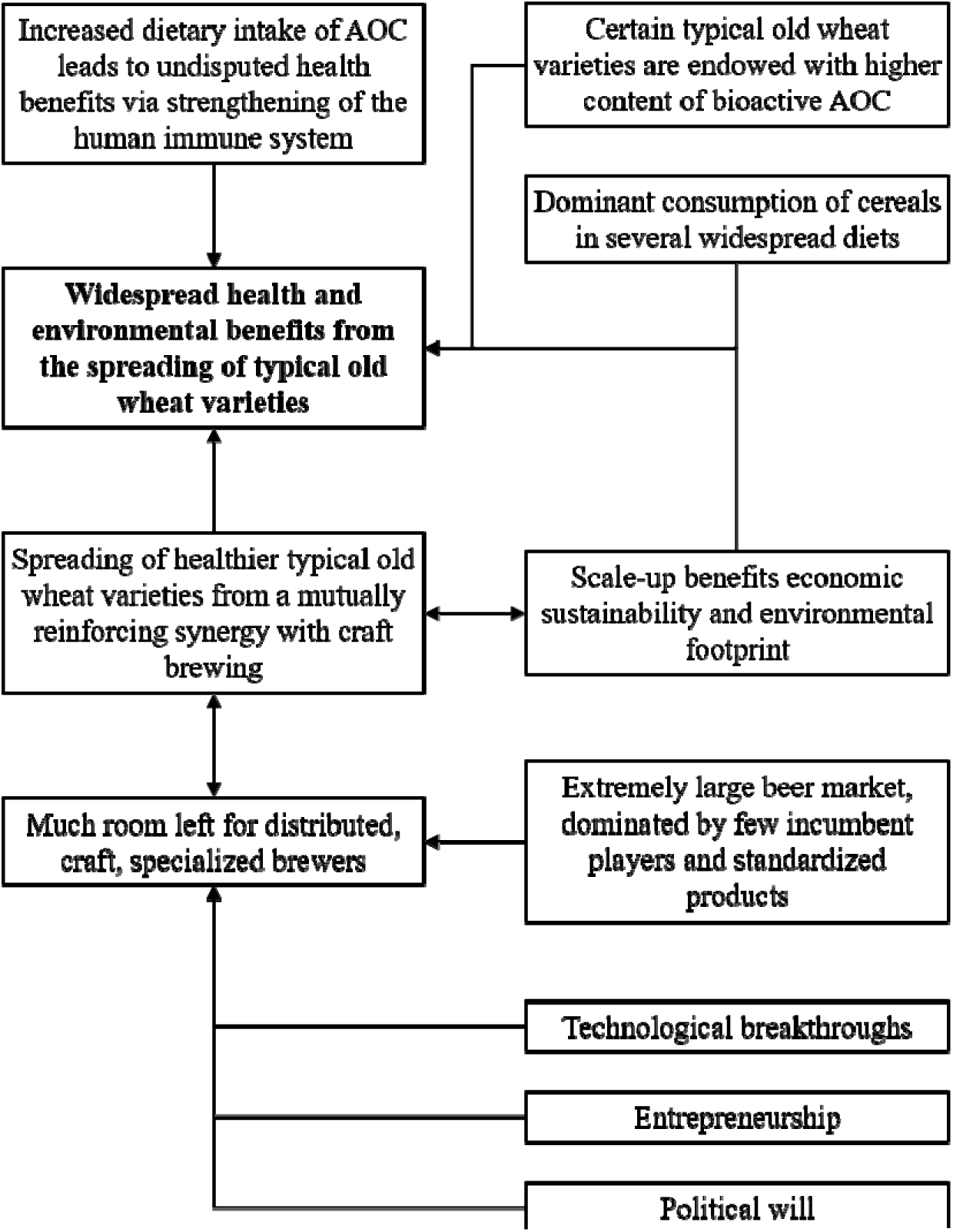
Simplified scheme of the vision developed in this study.

## 6 Conclusions

The synergy between beer-brewing by means of the more efficient novel technology described in Section 3, and the cultivation of typical old wheat varieties, whose exploitation in brewing was shown feasible in Section 4, appears as a viable pathway to improving public health as well as easing costs and environmental burdens of both the agricultural and health sectors.

As illustrated in Section 2, health benefits from increasing the dietary intake of antioxidant compounds, mostly in the form of flavonoids and other polyphenols of which certain old wheat varieties are especially endowed, are now undisputed, despite metabolic pathways conductive to the strengthening of the human immune system are still partially unknown (del Cornò et al., 2016; Pérez-Cano and Castell, 2016). Unfortunately, to the best knowledge of the authors, so far the public health component, along with the respective direct costs and environmental impact, has not been comprehensively accounted for in LCA studies of specific agricultural sectors, real or potential widespread shifts from certain crops to others, or changes in food consumption patterns related to the above-mentioned structural agricultural shifts. Proper consideration of the health issues connected to structural changes in crops and related foodstuff is recommended for future research.

On the contrary, recalling Section 1, comparative LCA studies of low-input and modern conventional cereal farming are widely available, converging towards contrasting patterns. In summary, lower yields often offset the easing of environmental burdens due to low-input farming, as well as the niche scale of low-input farming (among which most typical old wheat varieties) impairs the environmental footprint of distribution (Kulak et al., 2015).

However, yields of typical old wheat varieties has kept increasing (Ghiselli et al., 2010), as well as much room is left for further improvements (Ali et al., 2015). Perhaps even more important, the impressive extension of abandoned agricultural area, as mentioned in Section 2 for Italy and typical of most countries on the northern shores of Mediterranean, apart from the fraction left to urbanization, is likely to be most made up of less fertile soil, unsuitable for modern wheat varieties but suitable to grow typical old wheat varieties. Summarizing, the opportunity exists for a widespread transition back to old crops as well as for the respective yield increase, with the important result of their emergence from the niche to a real market scale.

As discussed in Section 5, confirming and even enhancing previously found results (Albanese et al., 2017), HC-assisted beer brewing, because of its competitive advantages, appears as a viable candidate to boost the spreading of healthier typical old wheat varieties by means of a purely market pathway. This opportunity could help easing the proposed taxation burden on the consumer side and avoid any regressive effect (García-Muros et al., 2017).

While further research is needed to optimize the HC-assisted brewing processes using different proportions of raw unmalted old wheat grains, a policy recommendation can be made to maintain tax benefits to agricultural craft breweries, allowing their recently observed strong rise to proceed at an accelerated pace (Fastigi et al., 2016). Such benefits could be even expanded for agricultural craft breweries based at organic or low-input farms growing typical old wheat varieties, especially those varieties endowed with the healthiest nutraceutical compounds.

The advocated paradigm shift from centralized and standardized production of commercial beer to widespread distributed production of craft beer recalls the basic aspects of the ongoing solar energy revolution (Meneguzzo et al., 2015). Both involve quite large markets and incumbent players constituting *de facto* oligopolies. In both cases, the actual feasibility of the production paradigm shift is closely tied to technology advancement: with solar energy, it occurred with the drastic price drop of solar photovoltaic modules, with old healthy wheat varieties and respective brewing it could occur as a consequence of the breakthrough technology proposed in this and previous studies by the authors (Albanese et al., 2017). Entrepreneurship attitude is crucial in both fields: tenths of thousand new companies and jobs in the millions were created in the solar energy business worldwide, while farmers and craft brewers are moving promising early steps also in search for a distinctive marketing message. Last, political will: as it was key to boost solar energy with feed-in tariffs and, later, auctions and tax relief, a daring fiscal leverage could boost both cultivation of typical old wheat varieties and brewing with the respective grains in the interest of public health and environment.

## Abbreviations

AOC: antioxidant compounds
ARG: antibiotic-resistant genes
BIAB: brew in the bag
CN: cavitation number
EBC: European Brewery Convention
EU: European Union
GHG: greenhouse gases
HC: hydrodynamic cavitation
LCA: Life Cycle Assessment
SG: specific gravity
SRM: Standard Reference Method

## Acknowledgements

This article is dedicated to Dr. Antonio Raschi (IBIMET-CNR), for all he has done in laying the foundations and pursuing the sustainable rural development. L.A. and F.M. were partially funded by Tuscany regional Government under the project T.I.L.A. (Innovative Technology for Liquid Foods, Grant N°. 0001276 signed on April 30, 2014). The research was carried out under a cooperation between CNR-IBIMET and the company Bysea S.r.l., with joint patent submitted on August 9, 2016, international application No. PCT/IT/2016/000194 “A method and relative apparatus for the production of beer”, pending. V. Marchioni (Melograno Società Cooperativa) is gratefully acknowledged for his continuous support and advice throughout this study. R. Lagalla (University of Palermo) is gratefully acknowledged for his invaluable support in the transfer of the patented brewing technology to industry and market.

